# Cell-derived plasma membrane vesicles are permeable to hydrophilic macromolecules

**DOI:** 10.1101/731364

**Authors:** AD Skinkle, I Levental

## Abstract

Giant Plasma Membrane Vesicles (GPMVs) are a widely used model system for biochemical and biophysical analysis of the isolated mammalian plasma membrane (PM). A core advantage of these vesicles is that they maintain the native lipid and protein diversity of the plasma membrane while affording the experimental flexibility of synthetic giant vesicles. In addition to fundamental investigations of PM structure and composition, GPMVs have been used to evaluate the binding of proteins and small molecules to cell-derived membranes, and the permeation of drug-like molecules through them. An important assumption of such experiments is that GPMVs are sealed; i.e. that permeation occurs by diffusion through the hydrophobic core rather than through hydrophilic pores. Here we demonstrate that this assumption is often incorrect. We find that most GPMVs isolated using standard preparations are passively permeable to various hydrophilic solutes as large as 40 kDa, in contrast to synthetic giant unilamellar vesicles (GUVs). We attribute this leakiness to relatively large and heterogeneous pores formed by rupture of vesicles from cells. These pores are stable and persist throughout experimentally relevant time scales. Finally, we identify preparation conditions that minimize poration and allow evaluation of sealed GPMVs. These unexpected observations of GPMV poration are of critical importance for interpreting experiments utilizing GPMVs as plasma membrane models, particularly for drug permeation and membrane asymmetry.

**STATEMENT OF SIGNIFICANCE:** A critical assumption in using Giant Plasma Membrane Vesicles to study membrane penetration and interactions is that these vesicles maintain the permeability barrier of the native membrane from which they form. Using large fluorescently-labeled hydrophilic probes, we demonstrate that this assumption is often incorrect and conclude that macromolecular solutes permeate GPMVs through stable pores formed during shear-induced rupture of vesicles from cells. Using these insights into the mechanisms of poration, we demonstrate an approach to isolate sealed GPMVs.

## INTRODUCTION

Model membranes are among the most widely used tools to investigate the physicochemical behaviors of biological lipids, interactions between lipids and proteins, and the properties of lipid assemblies. While synthetic lipid membranes effectively recapitulate certain aspects of biological membranes, it is not experimentally feasible to construct model systems with the appropriate lipid diversity and protein density of living cells. In this context, an important development in membrane biophysics has been the discovery (1) and characterization of Giant Plasma Membrane Vesicles (GPMVs), which are isolated from live mammalian cells as intact PM vesicles (2). These GPMVs fill a unique niche, maintaining the approximate lipid and protein content of living membranes while offering the experimental tractability of synthetic models (3). This combination has allowed investigation of PM composition (4), physical properties (5–7), dynamics (8, 9), and potentially signaling and transport (8, 10–13) in isolation from the complexity of living cells.

Because of these advantages, and despite several important caveats distinguishing isolated vesicles from living membranes (3), GPMVs have become an important and widely used tool for membrane biophysics. Perhaps their most notable feature is their spontaneous separation into coexisting liquid-ordered and disordered membrane phases, allowing investigations into the properties and compositions of ordered membrane domains, as models for lipid rafts (14–21). These vesicles have also been used in combination with mass spectrometric lipidomics to characterize the detailed and comprehensive lipid profiles of PMs and their remodeling during various cellular processes (22–24). Recently, GPMVs have also been employed to study mechanisms of viral binding and fusion to the PM (25), as well as toxin binding (26).

In addition to their considerable utility for basic research, GPMVs have recently gained attention for possible translational applications, particularly for their potential as drug delivery vehicles (27, 28) or models for studies of membrane permeability and poration (29–31). An important premise for these and other similar studies is that GPMVs are sealed to extravesicular compounds. This assumption stems from analogy to synthetic Giant Unilamellar Vesicles (GUVs), which are unambiguously sealed to permeation of hydrophilic solutes (26, 32). Here, we report the surprising finding that GPMVs are permeable to large, hydrophilic macromolecules. Probes as large as 40 kDa rapidly accumulate inside GPMVs, apparently through stable aqueous pores. We investigate the mechanism for this surprising permeability and suggest that stable pores are formed during shear-induced rupture of GPMVs from cells. Based upon these observations, we develop a technique to limit pore formation and image sealed GPMVs.

## MATERIALS AND METHODS

### Materials

The following materials were purchased for this study: Fast DiI (Life Technologies, Carlsbad, CA), fluorescein- and TMR-labeled dextrans (Sigma, St. Louis, MO), AlexaFluor-647-ATP (Fisher Scientific, Hampton, NH). The following lipids were purchased from Avanti Polar Lipids, Alabaster, AL: dipalmitoyl phosphatidylcholine (DPPC), dioleoyl PC (DOPC) and cholesterol.

### Cell culture

Rat basophilic leukemia (RBL) cells were maintained in medium containing 60% modified Eagle’s medium (MEM), 30% RPMI, 10% fetal calf serum, 2 mM glutamine, 100 units/mL penicillin, and 100 mg/mL streptomycin at 37°C in humidified 5% CO_2_.

### GPMV isolation and imaging

GPMVs were prepared as previously described (2). Briefly, cells were washed in GPMV buffer (10 mM HEPES, 150 mM NaCl, 2 mM CaCl2, pH 7.4) and then incubated with GPMV buffer supplemented with 25 mM paraformaldehyde (PFA) and 2 mM dithiothreitol (DTT) (or 2 mM N-ethylmaleimide (NEM) as indicated) for 1 h at 37°C (unless otherwise indicated). Before isolation, cell membranes were labeled with 5 μg/mL of the membrane marker FAST DiI for 10 min on ice. GPMVs were collected by sedimentation on ice, then imaged via confocal microscopy on a Nikon A1R.

### Quantification of GPMV leakiness

GPMVs were either prepared with (or incubated after isolation with) 250 nM AlexaFluor-647-ATP, 250 ng/mL fluorescein-labeled 3 kDa dextran. Confocal images were then taken with the fluorescent molecules still in solution. In these images, the lumens of “sealed” vesicles appeared dark relative to the background fluorescence, whereas leaky vesicles had bright fluorescent lumens (see Fig. 1). These images were analyzed using a custom-built algorithm in the open-source software Cell Profiler. GPMVs were identified and masked as circular objects then the median fluorescence intensity (to eliminate edge-effects) of the vesicle lumen was calculated for each vesicle. The fluorescence intensity of the bulk/background was calculated as the median intensity of all pixels outside the GPMVs. The “relative vesicle intensity” (i.e. the intensity of the vesicle lumen relative to the bulk) was calculated as: *relative vesicle intensity* = *I*_*vesicle*_ / *I*_*bulk*_

**Figure 1.**
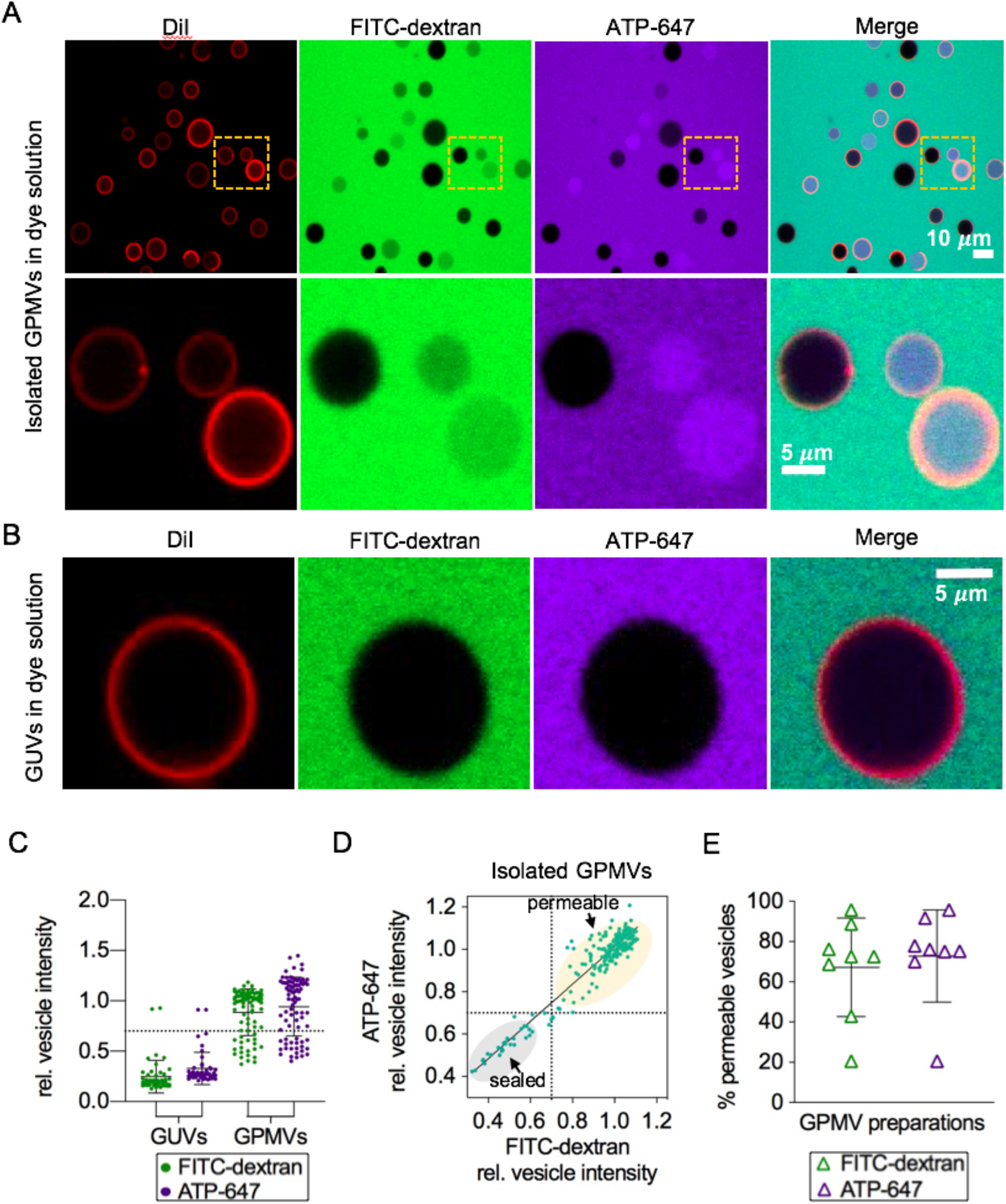
Large solutes permeate GPMVs. (A) Isolated GPMVs exposed to external FITC-modified 3 kDa dextran (FITC-dextran) and AlexaFluor-647 modified ATP analog (ATP-647) show intravesicular accumulation of probes in some vesicles. (B) In contrast, GUVs are not permeable to the same probes. This effect is quantified in (C) where points represent relative intensities (compared to bulk) of individual vesicles in a given preparation. A large population of permeable GPMVs (defined by a relative intensity > 0.7; see methods) is observed while almost all GUVs remain sealed. (D) Permeable GPMVs fill with both probes proportionally, revealing clear populations comprising sealed and permeable vesicles. (E) While the percentage of permeable vesicles varies considerably between preparations, most isolated GPMVs are permeable. Each point represents the percentage of permeable vesicles in a given preparation (N>100 vesicles / preparation).

Although most “leaky” vesicles had an internal fluorescence that was approximately equal to bulk fluorescence, some were slightly dimmer while others slightly brighter than the bulk fluorescence. Thus, a “leaky” vesicle was defined as one whose intensity was at least 70% of bulk fluorescence (relative vesicle intensity >0.7). Although this value is somewhat arbitrary, it was chosen because it consistently and clearly segregated the two populations of GPMVs that emerged in our analyses (see Fig. 1). Sealed vesicles also had non-zero “relative vesicle intensities”, which would appear to suggest that some permeation still occurred. Although we cannot rule this out, we believe this non-zero lumenal signal comes from analyzing images with a very bright fluorescent background. Even in confocal mode, some out-of-focus signal is detectable from above and below the vesicle, and this depends on the size of the vesicle. Thus, there are no truly “dark” pixels in our images, preventing the possibility of accurate background subtraction. An example of this effect (nonzero normalized intensity in putatively sealed vesicles) is shown in Fig. S1. Because of the large contrast between the “sealed vesicle” lumens and the bright bulk fluorescence, and because most putatively sealed vesicles had similar lumenal intensities, we believe there was minimal permeation of the fluorescent solutes into these.

### Preparation of giant unilamellar vesicles

GUVs were prepared by electroformation, as previously described (33). Briefly, a lipid mixture of dipalmitoyl phosphatidylcholine (DPPC), dioleoyl PC (DOPC) and cholesterol was prepared in chloroform with a molar ratio of 5:3:2 (lipids purchased from Avanti). The lipid mixture was applied to Pt-wires of a custom electroformation chamber and solvents were evaporated in a vacuum chamber. The electroformation chamber was filled with sucrose solution (0.1 M) and heated to 53°C. An alternating sinusoidal current was applied across the cell unit with 2.5 V, 10 Hz at 53 °C for 2 hours. GUVs were then imaged and analyzed as described above for GPMVs.

## RESULTS AND DISCUSSION

### GPMVs are permeable to hydrophilic macromolecules

GPMVs are widely used models of cellular plasma membranes and as such have been presumed to retain the barrier function of the native PM. This assumption has been reinforced by GPMVs’ superficial resemblance to synthetic GUVs, which generally do not have large hydrophilic pores. Surprisingly, we observed that GPMVs exhibited considerable heterogeneity in their permeability to extravesicular solutes in the form of fluorescently-labeled hydrophilic probes. When GPMVs were prepared in the presence of an Alexa647-modified ATP analog (2 kDa) or FITC-modified dextran (3 kDa), a major fraction of vesicles showed accumulation of these probes in the intravesicular space (Fig. 1A), a phenomenon hereafter referred to as permeation.

While fluorescent dextran was generally slightly less concentrated in the lumen of permeable GPMVs compared to the bulk extravesicular solution, fluorescent ATP, which can presumably interact with intravesicular proteins, was often clearly enriched (Fig. 1A inset). In striking contrast, synthetic giant unilamellar vesicles (GUVs) prepared by electroformation were almost uniformly impermeable to the same solutes, as exemplified in Fig. 1B and quantified in Fig. 1C. Within any given GPMV population, there was considerable variation in the accumulation of probes inside the vesicle, ranging from impermeable to slightly brighter than the bulk fluorescence of the extravesicular solution (Fig. 1D). Notably, all permeable GPMVs were similarly permeable to both fluorescent solutes, while GPMVs that were impermeable to dextran were also impermeable to fluorescent ATP (Fig. 1D). These observations allowed us to separate the GPMVs in a given preparation into clearly distinct sealed and permeable populations (Fig. 1D).

The fraction of permeable GPMVs was somewhat variable between repeated preparations (the reasons underlying this variability are discussed below); however, on average, the majority of vesicles were permeable to both fluorescent solutes (Fig. 1E). GPMVs were similarly permeable to FITC-modified dextran in the absence of fluorescent ATP (Fig. S2) revealing that permeation is not driven by the ATP analogue. These observations were not dependent on the chemicals chosen for GPMV production, as GPMVs prepared with N-ethylmaleimide were equivalently permeable to those prepared with PFA+DTT (Fig. S3)

Time-lapse imaging revealed that permeation occurred abruptly, and after the formation of an initially sealed vesicle. Fig. 2A and Video S1 show the temporal evolution of one such vesicle, which is sealed at an arbitrary starting point (approximately 30 mins after induction of vesiculation), but then rapidly fills with both macromolecular solutes within approximately one minute of the permeation-inducing event (the possible nature of the permeation-inducing event is discussed below). This permeation was not associated with any change in the gross morphology of the vesicle or its membrane, as shown by the lipid dye DiI (Fig. 2; bottom row). The kinetics of filling can be fit to a first-order rate constant, suggesting that both solutes reach our defined ‘permeability threshold’ (rel. vesicle intensity of 0.7) in <1 min. This rapid accumulation is inconsistent with the permeation of large hydrophilic probes through an intact membrane hydrophobic core, suggesting alternative permeation pathways. To test the mechanisms underlying these surprising observations, we evaluated the size, temperature, time, and mechanical perturbation dependence of GPMV permeability.

**Figure 2.**
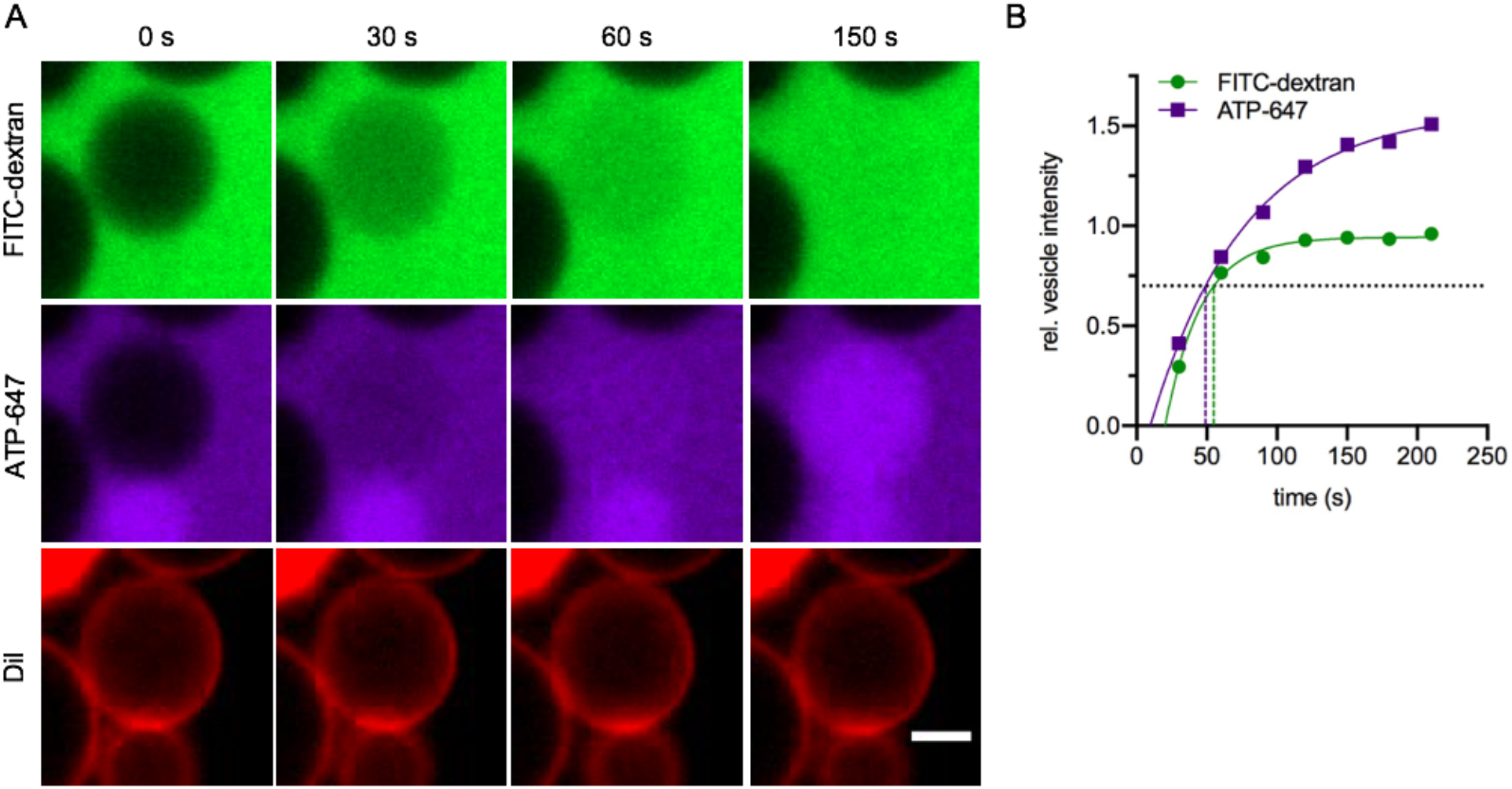
Large solutes permeate GPMVs rapidly. (A) Time-lapse imaging of GPMVs exposed to hydrophilic probes revealed vesicles fill with solutes on the order of one minute. (B) Dotted lines indicate time to vesicle filling (reaching a relative vesicle intensity of 0.7) from fits to first-order reaction kinetics. For this vesicle, the ‘filling time’ is ~55 s for FITC-dextran and ~49 s for ATP-647. Scale bar is 5 *μ*m.

### Large solutes permeate GPMVs via stable pores

To determine whether permeation through the GPMV membrane was occurring via well-defined pores, we tested permeation to fluorescent dextrans of increasing molecular weight. We observed a clear solute size dependence, with larger substrates permeating less efficiently. This dependence is clearly demonstrated by incubating a vesicle preparation with two differently sized dextran molecules labeled with two different fluorophores. As shown in Fig. 3A, some vesicles are completely sealed (dark in both channels), while a large fraction are completely permeable to 3 kDa dextran but not 60 kDa dextran (red only). This dependence of permeation on size of the solute was observed up to ~40 kDa, with decreasing efficiency for larger substrates (Fig. 3B). However, this size-dependence was not as sharp as might be expected for very consistent, homogeneous pores, suggesting that a distribution of pore sizes exists in any given population of vesicles.

**Figure 3.**
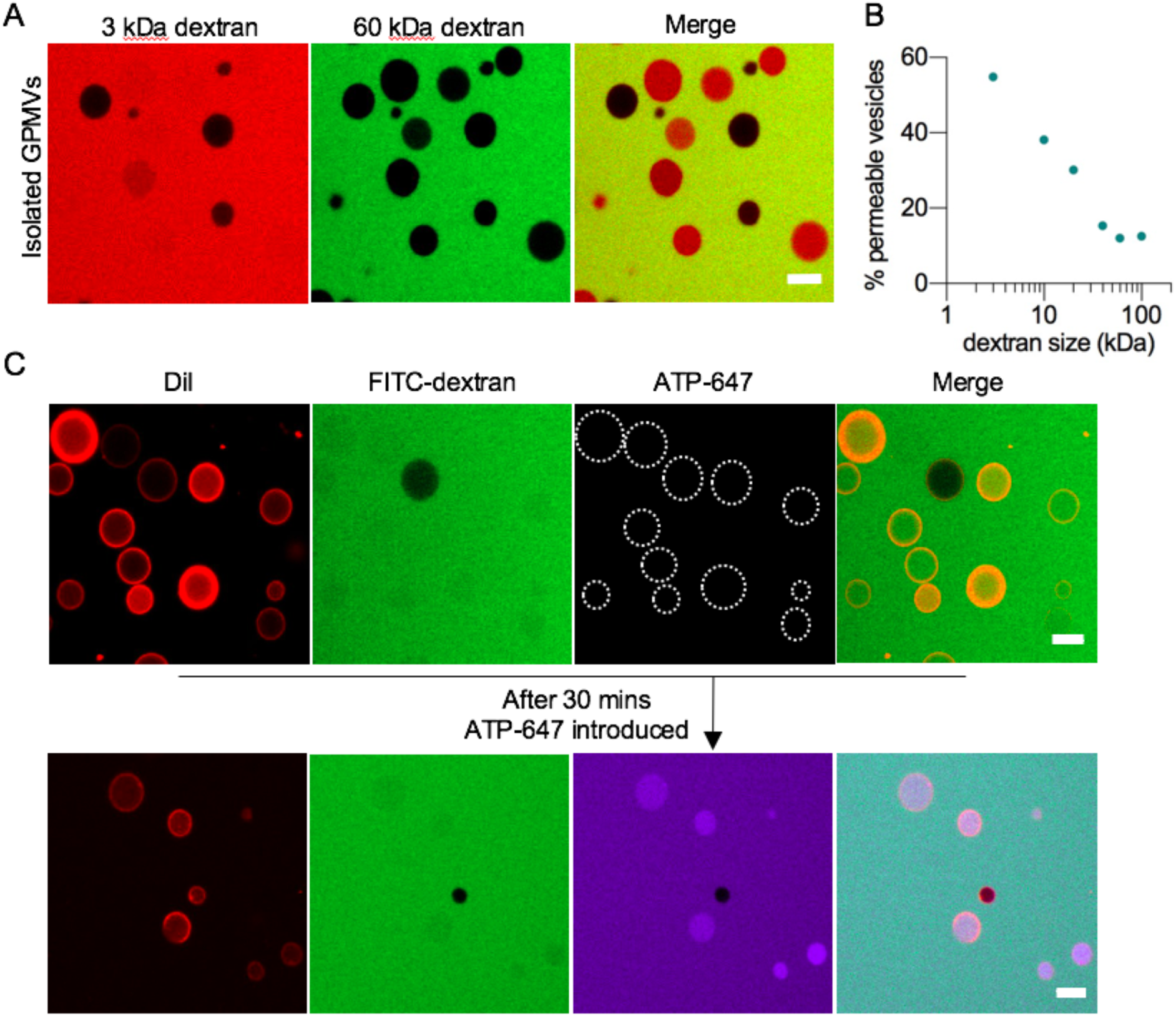
Large solutes permeate GPMVs via stable pores. When exposed to different molecular weight fluorescent dextrans, GPMV permeation showed a clear size dependence. (A) Vesicles that were permeable to 3 kDa dextran nearly completely excluded 60 kDa dextran. Higher molecular weight dextrans permeated GPMVs with decreasing efficiency up to 60 kDa as quantified in (B), suggesting the existence of pores of heterogeneous sizes. Each data point represents a population of GPMVs from the same preparation. (C) Vesicles that were prepared in the presence of FITC-dextran only, were exposed to ATP-647 30 mins after their isolation. Vesicles that were permeable to FITC-dextran at isolation were also permeable to ATP-647 after 30 mins suggesting pores remain intact at the vesicle membrane for extended periods of time. Scale bars are 10 *μ*m.

Finally, the possibility of stable pores was supported by the observations that permeable GPMVs retained their permeability well after isolation. GPMVs that were isolated in the presence of FITC-dextran alone were incubated for 30 mins at 37°C, then exposed to fluorescent ATP. After this incubation, vesicles that were permeable to one substrate were also permeable to the other, whereas vesicles that remained impermeable to dextran after 30 mins were also sealed to the second substrate (Fig. 3C). These observations suggest that stable pores (some of which can accommodate substrates as large as 40 kDa) are responsible for accumulation of solutes within GPMVs.

### GPMVs are not permeable immediately upon generation

To elucidate the conditions under which GPMVs are permeable, we first examined whether GPMVs were permeable immediately upon their generation or became permeable due to handling during/after generation. GPMV production was imaged from its initiation allowing observation of vesicles as they began to bud, inflated, then ultimately detached from cells. In these observations, both budding vesicles and almost all detached vesicles remained sealed to the small molecule probes (Fig. 4A), revealing that GPMVs are not initially permeable but become leaky after their generation.

**Figure 4.**
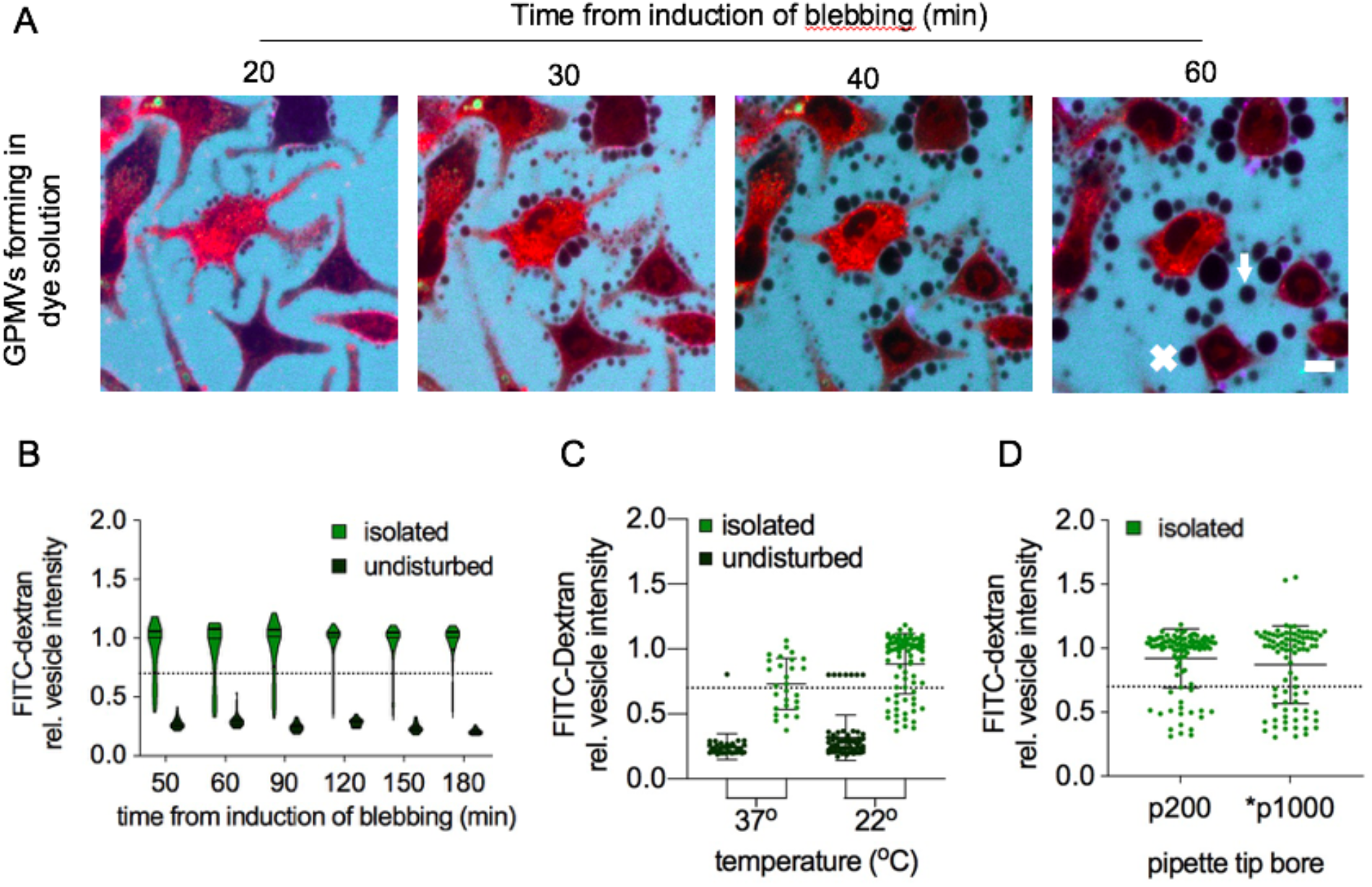
GPMVs become permeable upon isolation independent of temperature cycling and pipetting stress. (A) Time-lapse of cells forming GPMVs indicate GPMVs are not permeable upon their generation. Both blebbing (x) and detached (arrow) vesicles remained impermeable to both fluorescent probes. (B) GPMVs that had been isolated from cells were compared to those that were left undisturbed alongside the cells they formed from over the course of 2.5 hours. While undisturbed GPMVs remained impermeable, the isolated sample had a distribution of permeable and impermeable vesicles that remained consistent throughout time. (C) Keeping temperature constant between formation and isolation by isolating GPMVs at 37°C as opposed to 22°C did not reverse the permeation seen upon isolation. (D) Decreasing the shear strain of pipetting by increasing the bore of the pipette tip used for isolation did not diminish the population of permeable vesicles suggesting pipetting is not responsible for eliciting permeation. Asterisk indicates pipette tip was cut to increase bore. All results (B-D) repeated in the presence of ATP-647 as demonstrated in the supplement. Scale bar is 10 *μ*m.

To test the possibility that GPMVs become permeable over time regardless of handling, we observed GPMVs up to 2.5 hrs after isolation. Moreover, we compared GPMVs that were isolated from cells to those that were left undisturbed in the original plate in which they formed. Isolated GPMVs were characteristically heterogeneous in their accumulation of the extravesicular probes, with the majority being leaky to both solutes (Fig. 4B and S4A). Notably, the general distribution of permeable vesicles remained consistent throughout the time course, suggesting that the permeation-inducing even is not strongly time-dependent. In striking contrast, GPMVs that were not isolated from cells, but rather imaged in the same plate in which they were formed without manipulation, remained sealed throughout the time course, up to 3 h after induction of blebbing (Fig. 4B, S4A). These results suggested that the permeation-inducing event occurs during the process isolating GPMVs from the cells.

### Neither temperature changes nor shear from pipetting contribute to GPMV permeability

Given that GPMVs remain sealed until isolation from cells, we hypothesized that either temperature changes or the mechanical stress of pipetting during isolation was responsible for eliciting vesicle permeation. To test the effect of temperature changes, GPMVs were prepared at 37°C (as usual) then observed either at 37°C or 23°C. Both cases showed similar and characteristic distributions of permeable vesicles, while samples that were left undisturbed remained sealed regardless of temperature changes (Fig. 4C, S4B). Thus, temperature changes occurring during isolation are insufficient to elicit permeation. To test the possibility that mechanical stress of pipetting causes lipid bilayer disruptions large enough to allow permeation of solutes into the vesicle, GPMVs were isolated using pipette tips of varying bores and pipetted several times (Fig 4D, S4C). We also tested 1000 μL pipette tips from which the ends were cut to minimize shear stress on the vesicles. In all conditions, GPMVs displayed a characteristic distribution of permeable vesicles, suggesting shear during isolation had little effect on GPMV permeability.

### GPMVs become permeable upon rupture from cells

During the course of these investigations, we observed that plates of GPMV-forming cells that were mechanically jostled during preparation contained many fewer cell-attached GPMVs and many more permeable GPMVs. These observations suggested that shear from flow of the bulk solution surrounding GPMVs may be inducing permeability. To test this hypothesis directly, GPMV production was imaged while cells were immobilized on the microscope stage. As previously observed, unperturbed cells produced exclusively sealed GPMVs. Immediately after observation, the same plate of cells was jostled to induce flow of the bulk solution. After this perturbation, the previously sealed population of GPMVs now contained a subset of permeable vesicles (Fig. 5A).

**Figure 5.**
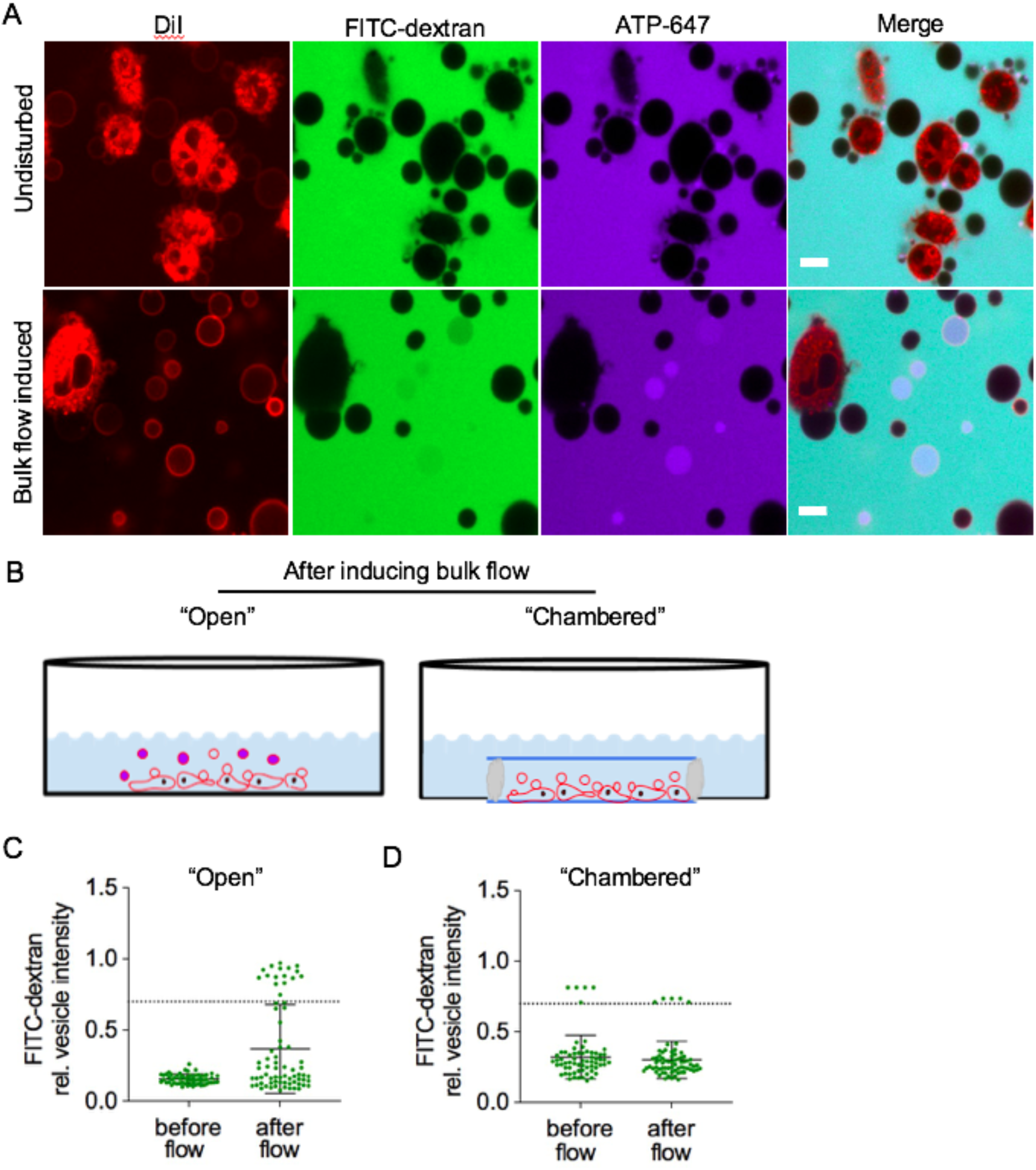
GPMVs become permeable upon shear-induced detachment from cells. (A) When a plate of cells forming GPMVs is moved, shear stress from movement of the bulk solution is induced and a previously sealed population of GPMVs becomes permeable. (B) GPMVs forming in this manner, named the “open” configuration become permeable after inducing shear, likely by traumatic rupture from cells. However GPMV formation can be induced in a sealed chamber which eliminates shear forces from movement of the bulk solution. Chambered GPMVs remain sealed when shearing conditions are induced. (C) When quantified, GPMVs forming in the “open” configuration became permeable upon generating shear from movement of the bulk solution. (D) While those GPMVs forming in the chambered configuration remain impermeable when the plate is moved. In (C-D) data points represent individual vesicles in a preparation of GPMVs. Scale bar is 10 *μ*m.

Given that pipetting of isolated GPMVs did not induce permeation but bulk fluid flow did, it is unlikely that shear forces from fluid flow damage intact isolated GPMVs. Rather, it seems more likely that shear forces result in a mechanical rupture of GPMVs from cells and it is this rupture event that results in the loss of bilayer integrity responsible for permeation. To confirm this inference, we compared the permeability of GPMVs produced in a standard “open” dish to a sealed chamber designed to prevent bulk fluid flow around the cells (Fig. 5B). In contrast to the “open” configuration where GPMVs became permeable upon inducing movement of bulk solution, GPMVs produced in the sealed chamber remained uniformly impermeable regardless of mechanical perturbation of the plate (Fig. 5D and S5B). When such sealed chambers were inverted, GPMVs separated from cells and settled to the bottom coverslip allowing them to be visualized separately from cells (Fig. 6A). Importantly, these GPMVs were almost exclusively sealed (Fig. 6B), offering an effective method to visualize sealed GPMVs.

**Figure 6.**
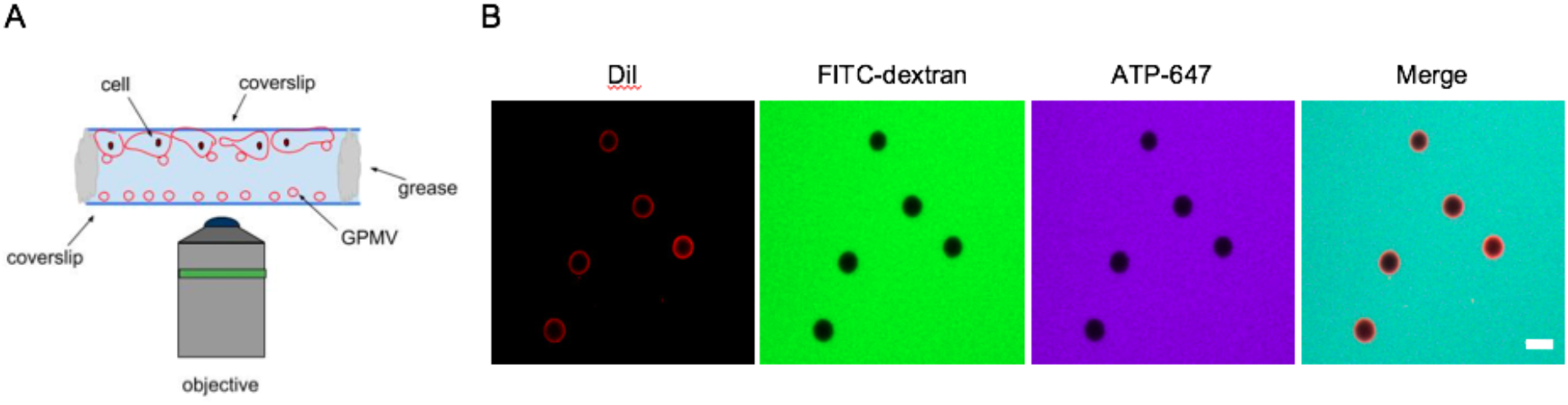
GPMVs formed in shear-protected conditions remain sealed. (A) GPMVs formed in a shear-protected chamber can be separated from cells by inverting the chamber. Because of their density, detached GPMVs sink towards the bottom coverslip allowing them to be visualized without also visualizing cells. (B) Importantly, these GPMVs remain sealed. Scale bar is 10 um.

## DISCUSSION

Here, we characterize the surprising observation that GPMVs, a widely used plasma membrane model, are passively permeable to macromolecular solutes. We show that GPMVs are permeable to solutes as large as 40 kDa and that they remain permeable after their isolation, regardless of the specific procedure used to isolate them. We observe a clear solute size-dependence, with the majority of vesicles being permeable to solutes below 10 kDa, and decreasing fractions being permeable to larger solutes up to a maximal cutoff of ~60 kDa. These observations combined suggest stable, somewhat variably sized, hydrophilic pores as the most likely potential mechanism underlying GPMV permeation.

Such pores are generally not observed in pure lipid model membranes, as confirmed here in GUVs (Fig. 1B-C), prompting the question of how they are formed and why they are stable in GPMVs. Our observations suggest a plausible model wherein shear stress induced by bulk fluid flow mechanically ruptures the GPMV away from the cell plasma membrane. This rupture may induce a membrane pore that can be stabilized by amphiphilic proteins in the GPMV lumen or unfolding of GPMV membrane proteins. Consistent with this inference, isolation of vesicles with no mechanical perturbation yields GPMVs that are almost completely sealed (Fig. 4).

How do some vesicles detach from the PM without forming a pore? One possibility involves protein machinery whose specific function is to mediate defect-free fission of membranes with the relevant toplogy, namely the endosomal sorting complexes required for transport (ESCRT) complex (34). We speculate that in some cells undergoing blebbing, ESCRT components assemble at the GPMV neck and catalyze scission, in analogy with the formation of intraluminal vesicles in late endosomes, microvesicles at the PM, or viruses budding from the PM (35). If this machinery is preempted by mechanical rupture, then a stable pore may form.

These findings are of particular relevance to studies using GPMVs for drug encapsulation (27, 28), therapeutic targeting (26, 27), and studies of membrane permeation (26, 29–32). Our suggestion for possible approaches to avoid leakage in GPMVs may be of value therein. Another important consideration is whether observations of protein binding to GPMVs may be biased by some probes/interactors leaking into the vesicle lumen.

## Author Contributions

AS and IL designed the studies and wrote the paper. AS performed all experimental work.

## Acknowledgements

All fluorescence microscopy was performed at the Center for Advanced Microscopy, Department of Integrative Biology & Pharmacology at McGovern Medical School, UTHealth. We gratefully acknowledge the members of the Levental lab for extensive discussions and support on this manuscript. Funding for this work was provided by the NIH/National Institute of General Medical Sciences (GM114282, GM124072, GM120351), the Volkswagen Foundation (grant 93091), and the Human Frontiers Science Program (RGP0059/2019). Authors have no competing interests.

## REFERENCES

1. Baumgart, T., A.T. Hammond, P. Sengupta, S.T. Hess, D.A. Holowka, B.A. Baird, and W.W. Webb. 2007. Large-scale fluid/fluid phase separation of proteins and lipids in giant plasma membrane vesicles. Proc. Natl. Acad. Sci. U. S. A. 104: 3165–3170.

2. Sezgin, E., H.-J. Kaiser, T. Baumgart, P. Schwille, K. Simons, and I. Levental. 2012. Elucidating membrane structure and protein behavior using giant plasma membrane vesicles. Nat. Protoc. 7: 1042–1051.

3. Levental, K.R., and I. Levental. 2015. Giant plasma membrane vesicles: models for understanding membrane organization. Curr. Top. Membr. 75: 25–57.

4. Sengupta, P., A. Hammond, D. Holowka, and B. Baird. 2008. Structural determinants for partitioning of lipids and proteins between coexisting fluid phases in giant plasma membrane vesicles. Biochim. Biophys. Acta. 1778: 20–32.

5. Kaiser, H.-J., H.-J. Kaiser, D. Lingwood, I. Levental, J.L. Sampaio, L. Kalvodova, L. Rajendran, and K. Simons. 2009. Order of lipid phases in model and plasma membranes. Proceedings of the National Academy of Sciences. 106: 16645–16650.

6. Veatch, S.L., P. Cicuta, P. Sengupta, A. Honerkamp-Smith, D. Holowka, and B. Baird. 2008. Critical fluctuations in plasma membrane vesicles. ACS Chem. Biol. 3: 287–293.

7. Moreno-Pescador, G., C.D. Florentsen, H. Østbye, S.L. Sønder, T.L. Boye, E.L. Veje, A.K. Sonne, S. Semsey, J. Nylandsted, R. Daniels, and P.M. Bendix. 2019. Curvature- and Phase-Induced Protein Sorting Quantified in Transfected Cell-Derived Giant Vesicles. ACS Nano. 13: 6689–6701.

8. Schneider, F., D. Waithe, M.P. Clausen, S. Galiani, T. Koller, G. Ozhan, C. Eggeling, and E. Sezgin. 2017. Diffusion of lipids and GPI-anchored proteins in actin-free plasma membrane vesicles measured by STED-FCS. Mol. Biol. Cell. 28: 1507–1518.

9. Worch, R., Z. Petrášek, P. Schwille, and T. Weidemann. 2017. Diffusion of Single-Pass Transmembrane Receptors: From the Plasma Membrane into Giant Liposomes. J. Membr. Biol. 250: 393–406.

10. Lewis, J.D., A.L. Caldara, S.E. Zimmer, S.N. Stahley, A. Seybold, N.L. Strong, A.S. Frangakis, I. Levental, J.K. Wahl 3rd, A.L. Mattheyses, T. Sasaki, K. Nakabayashi, K. Hata, Y. Matsubara, A. Ishida-Yamamoto, M. Amagai, A. Kubo, and A.P. Kowalczyk. 2019. The desmosome is a mesoscale lipid raft-like membrane domain. Mol. Biol. Cell. 30: 1390–1405.

11. Wei, X., H. Song, L. Yin, M.G. Rizzo, R. Sidhu, D.F. Covey, D.S. Ory, and C.F. Semenkovich. 2016. Fatty acid synthesis configures the plasma membrane for inflammation in diabetes. Nature. 539: 294–298.

12. Garten, M., L.D. Mosgaard, T. Bornschlögl, S. Dieudonné, P. Bassereau, and G.E.S. Toombes. 2017. Whole-GUV patch-clamping. Proceedings of the National Academy of Sciences. 114: 328–333.

13. Sedgwick, A., M. Olivia Balmert, and C. D’Souza-Schorey. 2018. The formation of giant plasma membrane vesicles enable new insights into the regulation of cholesterol efflux. Experimental Cell Research. 365: 194–207.

14. Lin, X., A.A. Gorfe, and I. Levental. 2018. Protein Partitioning into Ordered Membrane Domains: Insights from Simulations. Biophys. J. 114: 1936–1944.

15. Diaz-Rohrer, B.B., K.R. Levental, K. Simons, and I. Levental. 2014. Membrane raft association is a determinant of plasma membrane localization. Proc. Natl. Acad. Sci. U. S. A. 111: 8500–8505.

16. Sezgin, E., I. Levental, M. Grzybek, G. Schwarzmann, V. Mueller, A. Honigmann, V.N. Belov, C. Eggeling, U. Coskun, K. Simons, and P. Schwille. 2012. Partitioning, diffusion, and ligand binding of raft lipid analogs in model and cellular plasma membranes. Biochim. Biophys. Acta. 1818: 1777–1784.

17. Lorent, J.H., B. Diaz-Rohrer, X. Lin, K. Spring, A.A. Gorfe, K.R. Levental, and I. Levental. 2017. Structural determinants and functional consequences of protein affinity for membrane rafts. Nature Communications. 8.

18. Zhou, Y., K.N. Maxwell, E. Sezgin, M. Lu, H. Liang, J.F. Hancock, E.J. Dial, L.M. Lichtenberger, and I. Levental. 2013. Bile acids modulate signaling by functional perturbation of plasma membrane domains. J. Biol. Chem. 288: 35660–35670.

19. Gray, E., J. Karslake, B.B. Machta, and S.L. Veatch. 2013. Liquid general anesthetics lower critical temperatures in plasma membrane vesicles. Biophys. J. 105: 2751–2759.

20. Gray, E.M., G. Díaz-Vázquez, and S.L. Veatch. 2015. Growth Conditions and Cell Cycle Phase Modulate Phase Transition Temperatures in RBL-2H3 Derived Plasma Membrane Vesicles. PLoS One. 10: e0137741.

21. Johnson, S.A., B.M. Stinson, M.S. Go, L.M. Carmona, J.I. Reminick, X. Fang, and T. Baumgart. 2010. Temperature-dependent phase behavior and protein partitioning in giant plasma membrane vesicles. Biochim. Biophys. Acta. 1798: 1427–1435.

22. Levental, K.R., M.A. Surma, A.D. Skinkle, J.H. Lorent, Y. Zhou, C. Klose, J.T. Chang, J.F. Hancock, and I. Levental. 2017. ω-3 polyunsaturated fatty acids direct differentiation of the membrane phenotype in mesenchymal stem cells to potentiate osteogenesis. Sci Adv. 3: eaao1193.

23. Levental, K.R., J.H. Lorent, X. Lin, A.D. Skinkle, M.A. Surma, E.A. Stockenbojer, A.A. Gorfe, and I. Levental. 2016. Polyunsaturated Lipids Regulate Membrane Domain Stability by Tuning Membrane Order. Biophys. J. 110: 1800–1810.

24. Levental, K.R., E. Malmberg, Y.-Y. Fan, R. Chapkin, R. Ernst, and I. Levental. Homeostatic remodeling of mammalian membranes in response to dietary lipids is essential for cellular fitness.

25. Yang, S.-T., A.J.B. Kreutzberger, V. Kiessling, B.K. Ganser-Pornillos, J.M. White, and L.K. Tamm. 2017. HIV virions sense plasma membrane heterogeneity for cell entry. Sci Adv. 3: e1700338.

26. Manni, M.M., J. Sot, and F.M. Goñi. 2015. Interaction of Clostridium perfringens epsilon-toxin with biological and model membranes: A putative protein receptor in cells. Biochim. Biophys. Acta. 1848: 797–804.

27. Patel, J.M., V.F. Vartabedian, E.N. Bozeman, B.E. Caoyonan, S. Srivatsan, C.D. Pack, P. Dey, M.J. D’Souza, L. Yang, and P. Selvaraj. 2016. Plasma membrane vesicles decorated with glycolipid-anchored antigens and adjuvants via protein transfer as an antigen delivery platform for inhibition of tumor growth. Biomaterials. 74: 231–244.

28. Gadok, A.K., D.J. Busch, S. Ferrati, B. Li, H.D.C. Smyth, and J.C. Stachowiak. 2016. Connectosomes for Direct Molecular Delivery to the Cellular Cytoplasm. J. Am. Chem. Soc. 138: 12833–12840.

29. Pae, J., P. Säälik, L. Liivamägi, D. Lubenets, P. Arukuusk, Ü. Langel, and M. Pooga. 2014. Translocation of cell-penetrating peptides across the plasma membrane is controlled by cholesterol and microenvironment created by membranous proteins. J. Control. Release. 192: 103–113.

30. Säälik, P., A. Niinep, J. Pae, M. Hansen, D. Lubenets, Ü. Langel, and M. Pooga. 2011. Penetration without cells: membrane translocation of cell-penetrating peptides in the model giant plasma membrane vesicles. J. Control. Release. 153: 117–125.

31. Dubavik, A., E. Sezgin, V. Lesnyak, N. Gaponik, P. Schwille, and A. Eychmüller. 2012. Penetration of amphiphilic quantum dots through model and cellular plasma membranes. ACS Nano. 6: 2150–2156.

32. Faust, J.E., P.-Y. Yang, and H.W. Huang. 2017. Action of Antimicrobial Peptides on Bacterial and Lipid Membranes: A Direct Comparison. Biophys. J. 112: 1663–1672.

33. Dimitrov, D.S., and M.I. Angelova. 1988. Lipid swelling and liposome formation mediated by electric fields. Bioelectrochemistry and Bioenergetics. 19: 323–336.

34. Jimenez, A.J., P. Maiuri, J. Lafaurie-Janvore, S. Divoux, M. Piel, and F. Perez. 2014. ESCRT Machinery Is Required for Plasma Membrane Repair. Science. 343: 1247136–1247136.

35. McDonald, B., and J. Martin-Serrano. 2009. No strings attached: the ESCRT machinery in viral budding and cytokinesis. J. Cell Sci. 122: 2167–2177.

