## Supplementary Information for "Cell-derived plasma membrane vesicles are permeable to hydrophilic macromolecules"

### SUPPLEMENTAL INFORMATION

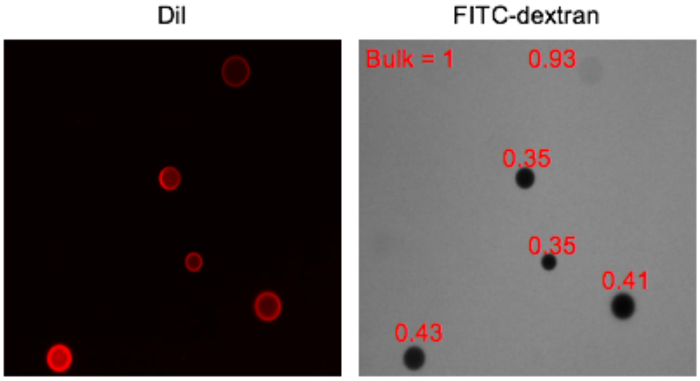


**Figure S1. Sealed vesicles have non-zero normalized intensities.** Out of focus light from planes above and below confocal slices contribute to a non-zero normalized intensity, even for sealed GPMVs. Normalized intensity values are written above their corresponding vesicle to demonstrate this effect.

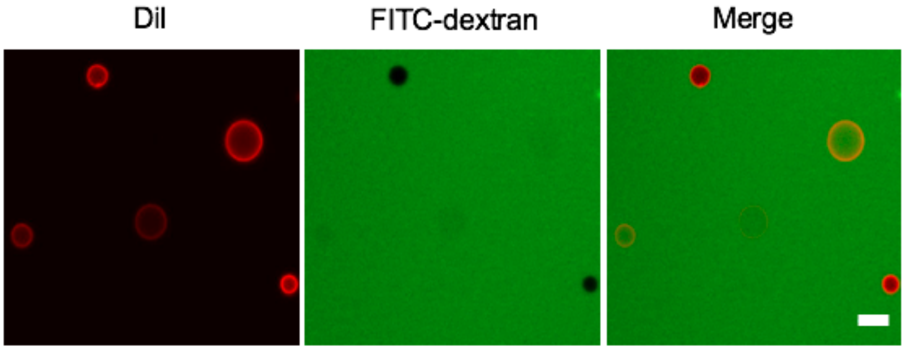


**Figure S2. GPMV permeation does not require ATP analogue.** GPMVs exposed to only FITC modified dextran (3kDa) still exhibited permeation demonstrating that addition of Alexa-fluor 647 modified ATP analogue is not necessary to elicit permeation. Scale bar is 10 $\mu$m.

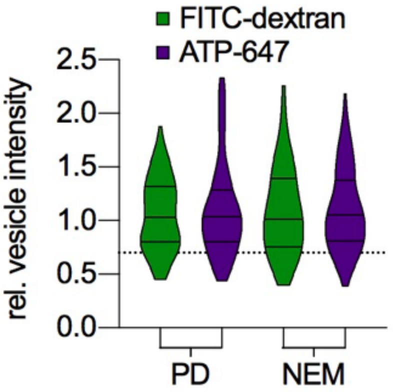


**Figure S3. Preparation reagents do not influence permeation.** GPMVs prepared by different vesiculation reagents, either with paraformaldehyde and dithiotreitol (PD) or with n-ethylmalemide (NEM), exhibited little difference in distribution of permeable GPMVs.

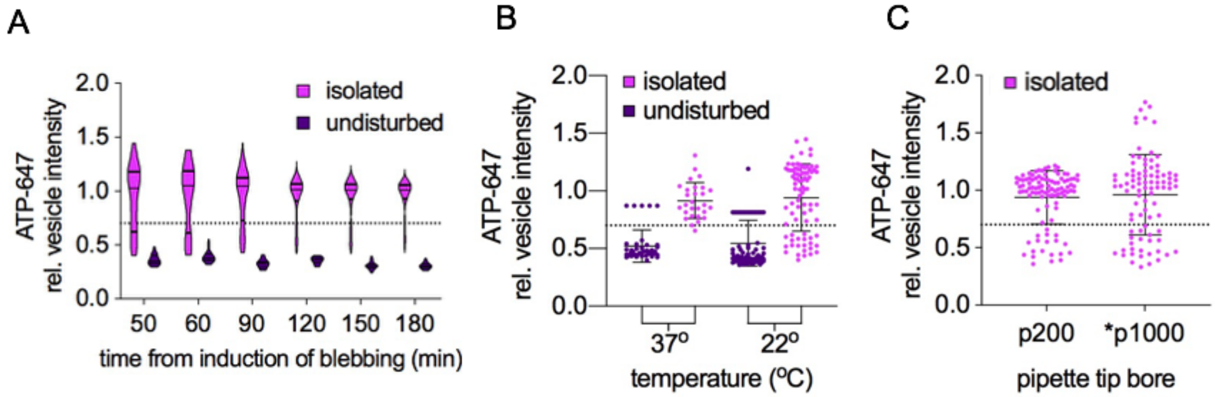


**Figure S4. GPMVs become permeable upon isolation independent of temperature cycling and pipetting stress: ATP-647.** (A-C) Correspond to the experiments from Fig 4 (B-D) but in the presence of ATP-647. (A) GPMVs that had been isolated from cells were compared to those that were left undisturbed alongside the cells they formed from over the course of 2.5 hours. While undisturbed GPMVs remained impermeable to ATP-647, the isolated sample had a distribution of permeable and impermeable vesicles that remained consistent throughout time. (B) Keeping temperature constant between formation and isolation by isolating GPMVs at 37°C as opposed to 22°C did not reverse the permeation seen upon isolation. (C) Decreasing the shear strain of pipetting by increasing the bore of the pipette tip used for isolation did not diminish the population of vesicles permeable to ATP-647 suggesting pipetting is not responsible for eliciting permeation. Asterisk indicates pipette tip was cut to increase bore.

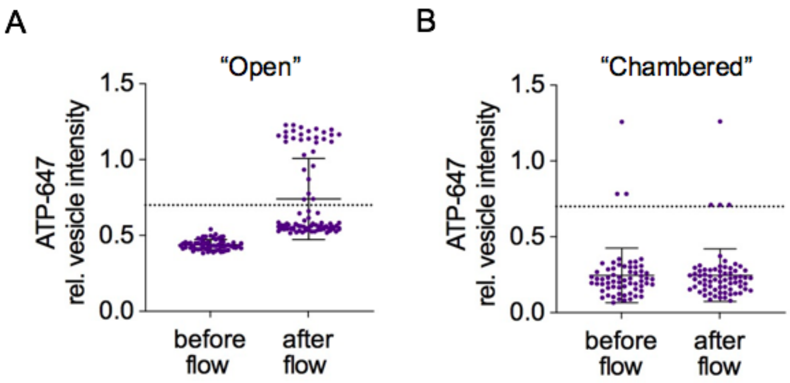


**Figure S5. GPMVs become permeable upon shear-induced detachment from cells: ATP-647.** (A-B) Correspond to experiments (C-D) from Fig. 5 but in the presence of ATP-647. (A) GPMVs forming in the "open" configuration were similarly permeable to ATP-647 when shearing from movement of the bulk solution was induced. (B) While those GPMVs forming in a "chambered" configuration remained sealed to ATP-647 under the same shear-inducing conditions.

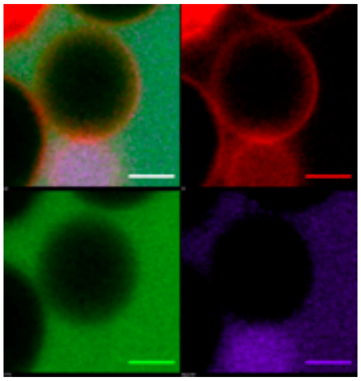

**Video S1. Time-lapse imaging of a GPMV becoming permeable.** A time-lapse was taken of GPMVs forming in the presence of FITC-dextran (green) and ATP-647 (purple). Membranes were labeled in FASTDiI (red). Video depicts solutes accumulating in the vesicle within several frames, as quantified in Fig. 2. Frames are 30s. Scale bar is 5 $\mu$m.
